# Natural variation in the maternal and zygotic mRNA complements of the early embryo in *Drosophila melanogaster*

**DOI:** 10.1101/2021.11.01.466760

**Authors:** Anna A. Feitzinger, Anthony Le, Ammon Thompson, Mehnoor Haseeb, Mohan K. Murugesan, Austin M. Tang, Susan E. Lott

## Abstract

Maternal gene products supplied to the egg during oogenesis drive the earliest events of development in all metazoans. After the initial stages of embryogenesis, maternal transcripts are degraded as zygotic transcription is activated, this is known as the maternal to zygotic transition (MZT). Altering the abundances of maternally deposited factors in the laboratory can have a dramatic effect on development, adult phenotypes and ultimately fitness. Zygotic transcription activation is a tightly regulated process, where the zygotic genome takes over control of development from the maternal genome, and is required for the viability of the organism. Recently, it has been shown that the expression of maternal and zygotic transcripts have evolved in the Drosophila genus over the course of 50 million years of evolution. However, the extent of natural variation of maternal and zygotic transcripts within a species has yet to be determined. We asked how the maternal and zygotic pools of mRNA vary within and between populations of *D. melanogaster*. In order to maximize sampling of genetic diversity, African lines of *D. melanogaster* originating from Zambia as well as DGRP lines originating from North America were chosen for transcriptomic analysis. Single embryo RNA-seq was performed at a stage before and a stage after zygotic genome activation in order to determine which transcripts are maternally deposited and which are zygotically expressed within and between these populations. Differential gene expression analysis has been used to quantify quantitative changes in RNA levels within populations as well as fixed expression differences between populations at both stages. Generally, we find that maternal transcripts are more highly conserved, and zygotic transcripts evolve at a higher rate. We find that there is more within population variation in transcript abundance than between populations and that expression variation is highest post-MZT between African lines. Determining the natural variation of gene expression surrounding the MZT in natural populations of *D. melanogaster* gives insight into the extent of how a tightly regulated process may vary within a species, the extent of developmental constraint at both stages and on both the maternal and zygotic genomes, and reveals expression changes allowing this species to adapt as it spread across the world.

## Introduction

Over the course of the development of multicellular organisms, an embryo that starts with a single nucleus undergoes divisions with dynamic changes in gene expression to give rise to a functional organism. This can require tight temporal and spatial control of gene expression throughout development, which is complicated by the fact that early development requires the coordination of gene expression across two different genomes. The earliest steps of embryonic development are under complete control of gene products supplied by the maternal genome before developmental control is transferred to the zygote (Tadros & Lipshitz, 2009, Vastenhouw et al., 2019). This process, where control of development is handed off between the maternal and zygotic genomes, is known as the maternal to zygotic transition (MZT) and has been the subject of study of many model organisms (Vastenhouw et al., 2019). In *Drosophila melanogaster*, maternal RNAs are transcribed during oogenesis in specialized cells called nurse cells and then supplied to the oocyte (Lasko, 2012). During the MZT, these maternal RNAs are degraded as the zygotic genome is activated, ~ 2.5 hours after fertilization (Bashirullah et al., 1999). Levels of many transcripts produced by both the maternal and zygotic genomes appear invariant across the MZT, indicating precise coordination of maternal degradation and zygotic transcription (Lott et al., 2011).

Given the importance of early development to organism survival and its dependence on precise regulation and coordination across the maternal and zygotic genomes, it may be unsurprising that a previous study found a high level of conservation of transcript levels across *Drosophila* species (Atallah & Lott, 2018). However, the same study (Atallah & Lott, 2018) was able to identify some changes in transcript representation and abundance across the 50 million years of divergence time of *Drosophila* at both the maternal and zygotic stages, and map these changes on the phylogeny. Given that these species have significant differences in the environments in which they develop, some of these changes may be functionally critical to developing under different conditions. Correlations of maternal and zygotic transcript levels decreased with evolutionary divergence, and changes in transcript representation were found even between closely related species (Atallah & Lott, 2018). But a significant question remains: do differences in maternal and zygotic transcript levels evolve in the comparatively short evolutionary timescales represented by different populations within a species? Understanding the extent of changes in transcript levels in these critical developmental stages of populations within a species can inform us about the timescale of evolutionary change. Exploring the types of genes that change in the context of different populations may also be a promising avenue for understanding the functions and potential adaptive value of these changes.

In this study, we sought to determine the extent of variation in maternal and zygotic embryonic transcriptomes between populations. To maximize the probability of observing differences, we chose populations of *D. melanogaster* that were likely to be highly genetically diverged. As a species, there is evidence that *D. melanogaster* has its origins in Sub-Saharan Africa (Begun & Aquadro, 1993; Pool & Aquadro, 2006). Approximately 10,000 years ago, it is likely that *D. melanogaster* began to expand beyond Sub-Saharan Africa (Li & Stephan 2006, Thornton & Andolfatto 2006) and eventually into northern Africa, Asia, and Europe. Only within the past few hundred years were North American populations of *D. melanogaster* founded (Lintner, 1882). With the expansion of *D.melanogaster* out of Sub-Saharan Africa, there was likely a significant loss in genetic diversity (Kauer et al., 2003). Efforts to sequence genomes from different lines and geographic populations of *D. melanogaster*, including African populations, has been ongoing in order to understand underlying genetic variation and the demographic history of the species (Pool & Aquadro, 2006). Taking advantage of the large number of sequenced genomes and RNA sequencing technology, it has more recently become possible to interrogate correlations between genetic variation and transcriptome diversity. For instance, transcriptomic variation of African populations of *D.melanogaster*, which have more genetic variation than European populations hasbeen characterized for adult flies (Hutter et al., 2008). This has brought to light the extent of differential gene expression between these populations within the same species. In order to determine the constraints on maternal and zygotic transcript levels within *D.melanogaster* we chose to sample lines from Sub-Saharan Africa where this species originated, and lines from North America.

Here, we address how the maternal and zygotic transcriptomes controlling the critical processes in early embryogenesis differ between populations of *D. melanogaster*. We performed RNA-Seq on embryos from four lines from Zambia and four lines from North America (DGRP lines), from two developmental stages, one stage where all transcripts present are maternal in origin and the other after zygotic genome activation. Transcript level variation was quantified within two populations as well as putative fixed differences in gene expression between them. We discovered that variation of both maternal and zygotic transcript levels is higher within populations than between populations. We find that there is more expression variation within the Zambia population at both stages relative to the Raleigh population. We observe an enrichment on the X chromosome for maternally deposited mRNAs that are differentially deposited between the two populations. Additionally, we find less transcript level variation between any two of our *D. melanogaster* lines than between species of *Drosophila*. Overall, our results demonstrate that expression level variation at these two stages is consistent with what is known about the differences in genetic variation between these populations. Furthermore, differences in transcript levels at these two stages between populations of *D. melanogaster* recapitulate what is known between species of *Drosophila*.

## Materials and Methods

### Embryo collection and sequencing library generation

Fly populations from Zambia (courtesy of the Langley Lab, University of California, Davis) and Raleigh, NC, USA (the DGRP lines; MacKay et al., 2012) were population controlled on cornmeal fly food at 25 degrees C. Four lines from Zambia (ZI050, ZI094, ZI160, ZI470) and four lines from Raleigh (RAL307, RAL357, RAL360, RAL517) were selected for embryo collection. Embryos were dechorionated using 50% bleach, and imaged on a Zeiss Axioimager, under halocarbon oil, to determine stage. Since embryos were collected from a large number of mothers, it is unlikely that multiple samples came from the same mother. Stage 2 and late stage 5 embryos were identified based on morphology. Stage 2 embryos were selected based on the vitelline membrane retracting from both the anterior and posterior poles, prior to when pole cells become visible. Late stage 5 embryos were chosen based on having completed cellularization, but not yet having started gastrulation. Embryos were then removed from the slide with a brush, cleaned of excess oil, placed into a drop of Trizol reagent (Ambion), and ruptured with a needle, then moved to a tube with more Trizol to be frozen at −80 ° C until extraction. RNA and DNA were extracted as in the manufacturer’s protocol, with the exception of extracting in an excess of reagent (1 mL was used) compared to expected mRNA and DNA concentration. Extracted DNA for stage 5 embryos was used for genotyping for sex as in Lott et al, 2011, XY embryos were selected for transcriptomic analysis, due to the incomplete nature of X chromosomal dosage compensation in XX embryos at this stage (Lott et al, 2011).

RNA-Seq libraries were prepared using poly-A enrichment for each of the 8 lines (4 Zambia lines and 4 Raleigh lines), for both stage 2 and stage 5, with 3 replicates each, for a total of 48 libraries. These samples were sequenced 100bp, paired-end, on an Illumina HiSeq4000. The sequencing was carried out by the DNA Technologies and Expression Analysis Core at the UC Davis Genome Center, supported by NIH Shared Instrumentation Grant 1S10OD010786-01.

### Data Processing

Reads were trimmed and adapters removed using Cutadapt (Martin 2011), and gently (PHRED Q < 20) trimmed for quality (MacManes, 2014). Mapping was done with the D. melanogaster Flybase genome release 6.18 and associated annotation file using hisat2 version 2.1.0 (Pertea et al., 2016) using default parameters. Gene level counts were generated using featureCounts of the subRead (Liao et al., 2014) package in R (R version 3.4.1, R Core Team, 2017). Counts were normalized to sequencing depth and RNA composition using DEseq2’s median of ratios. Count data can be found in Supplemental File 1.

### Data availability

All raw and processed data are available at NCBI/GEO under accession number X. Processed data (transcript level counts) is also available in Supplemental File 1.

### Hierarchical Clustering and PCA Analysis

We performed hierarchical clustering analysis in R using the hclust function. A dissimilarity matrix (*dist()*) of one minus the Spearman correlation (*cor()*) was used for hierarchical clustering. Principal component analysis (PCA) was also performed in R using the *prcomp()* function.

### Determining on or off State

To determine whether a gene was likely to be transcribed based on the count data, we ran Zigzag (Thompson et al., 2020) on our data. A full description of how this program was utilized, see Supplemental File 2.

### Differential Expression Analysis

Differential expression analysis was done using the DEseq2 (Love et al., 2014) package in R. Using DEseq2, we implemented the LRT (likelihood ratio test). For within-population analysis the replicates for each line were given the same label for the design matrix. For determining the differences between populations, we labeled lines as either Raleigh or Zambia in the design matrix and implemented the LRT test. When comparing the number of DE genes within and between populations, the number of DE genes is divided by the number of genes expressed in order to compare % DE genes between stages. We counted a gene as expressed in the total number of genes expressed for normalization if the gene was expressed in at least one line, as described above.

For pairwise differences between lines, DEseq2 was run on every possible combination of pairs. Since there are more between population pairs than within population pairs, we ran bootstrapping in R in order to compare the number of DE genes between lines of the same population and between lines of different populations. To test if the distributions of bootstrapped averages were significantly different from one another, we implemented a Wilcoxon rank sum test in R.

For differential expression analysis between species we used RNA-seq data previously generated in the lab (Atallah & Lott, 2018) from *D. simulans, D. sechellia, D. erecta*, and *D. yakuba*. Reads were aligned using HISAT2 followed by FeatureCounts to generate expression levels in counts. Counts were then normalized using the norm() function in DESeq2. Only genes which had orthologs in all seven species were considered. An expression cut off of 3 counts was used to determine which genes were considered expressed in each line.

### Test of Enrichment on Autosomes or Sex Chromosomes

To determine whether there wasenrichment of DE genes on either the autosomes or sex chromosomes the chromosomal location of each DE gene was determined. Number of DE genes per chromosome was normalized to the number of genes expressed on the chromosome. We implemented a Fisher’s exact test in R to determine if there is a significant difference in how many DE genes are on autosomes compared to the X chromosome. This was performed by doing individual tests between the number of DE genes on each autosome and the X.

### Heat Shock of Embryos

We adapted the heat shock and embryo survival protocols from (Lockwood et al., 2017). Flies aged 3-5 days were allowed to lay on a clearance plate for one hour. Plates were then swapped with clear agar collection plates with additional yeast and flies allowed to lay for an additional hour in order to collect 0-1 hour aged embryos. Plates were then wrapped in parafilm and fully submerged in a heat bath at the given temperature for 40 minutes. Embryos were then grouped in a line of 20 embryos using a brush. Proportion of embryos hatched was assayed 48 hours after heat shock to determine embryo survival. Three temperatures were assayed.

## Results

In order to investigate the natural variation of RNA levels within a species at stages of embryogenesis controlled by maternal and zygotic genomes, we sequenced embryonic transcriptomes from different *D. melanogaster* populations. Single embryos were collected at a stage in which all RNA has been maternally provided (Bownes’ stage 2; Bownes 1975), and another stage after zygotic genome activation (late stage 5; or end of blastoderm stage). In order to maximize genetic diversity, we chose four lines from Zambia and four Drosophila Genetics Reference Panel (DGRP) lines from Raleigh, North Carolina. Three biological replicates were sequenced per line and stage. An average of 2.83 and 2.89 million high-quality 100 bp paired-end reads were mapped to the *D. melanogaster* genome from the Zambia and Raleigh lines, respectively. Hierarchical clustering of the transcriptomes resulted in samples clustering initially by stage then by population, with the exception of one Raleigh line whose stage 5 sample fell outside the three other stage 5 Raleigh samples. When we included transcriptomes from an outgroup, *D.simulans*, which share a common ancestor ~2.5 MYA with *D. melanogaster* (McDermott & Kliman, 2008), to the clustering, the *D. simulans* samples clustered by stage with, but outside of, the *D. melanogaster* transcriptomes (Figure 1A). Principal component analysis also separates individual lines by stage with the corresponding principal component (PC1) representing nearly 80% of the variation (Figure 1B).

**Figure 1.**
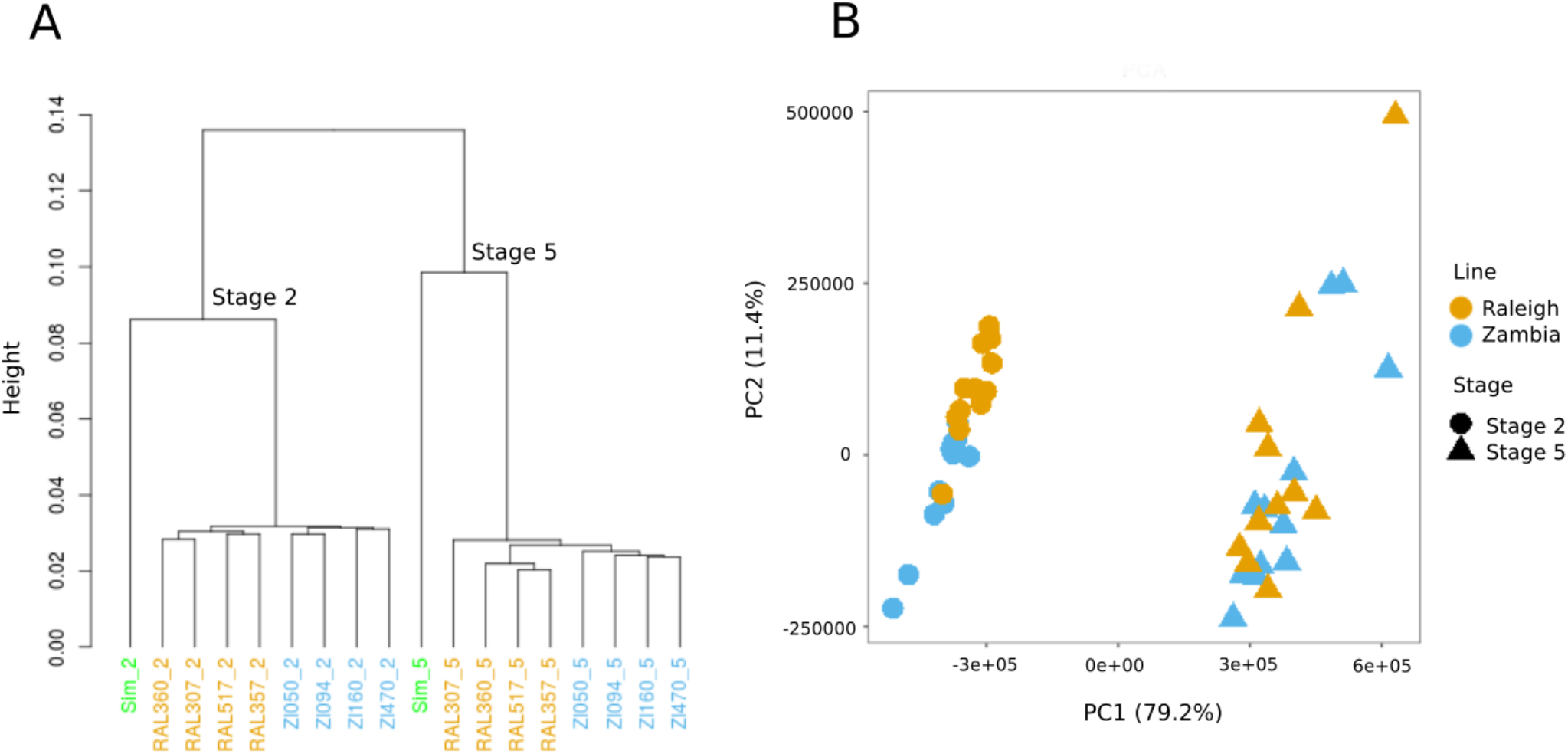
Populations are distinct at each developmental stage. A) Hierarchical clustering of transcriptomes from stage 2 (labels ending with _2) and stage 5 (labels ending with _5) embryos, from 8 lines of *D. melanogaster*, four from Raleigh (RAL, orange) and four from Zambia (ZI, blue), with closely related species *D. simulans* (Sim, in green text) as an outgroup. Samples cluster first by stage, then by species, then by population. B) PCA shows that these same samples separate first by stage (PC1, which explains a large proportion of the variance at 79.2%), then by population (PC2, 11.4% of the variance), though more distinctly at stage 2 than stage 5.

### Expression Variation Differs Within Populations

To explore the patterns of variation in the maternal and zygotic embryonic transcriptomes within and between populations of *D. melanogaster*, we performed differential expression (DE) analysis on our transcriptomic dataset. First, we asked how many genes are differentially expressed within each population, Zambia or Raleigh, at maternal and zygotic stages of development. To do this we implemented a likelihood ratio test in DESeq2. We normalized our differential expression results to numbers of genes expressed (see Methods) at each stage in order to compare proportions of genes differentially expressed between stages. We found that overall, there are more differentially expressed genes at stage 5 than at stage 2 within both populations (Fig 2A). This is consistent with previous findings between species that zygotic gene expression evolves faster than maternal gene expression (Atallah and Lott, 2018). Strikingly, there are many more differentially expressed genes at both stage 2 and stage 5 within the Zambia population than between Raleigh lines.

**Figure 2.**
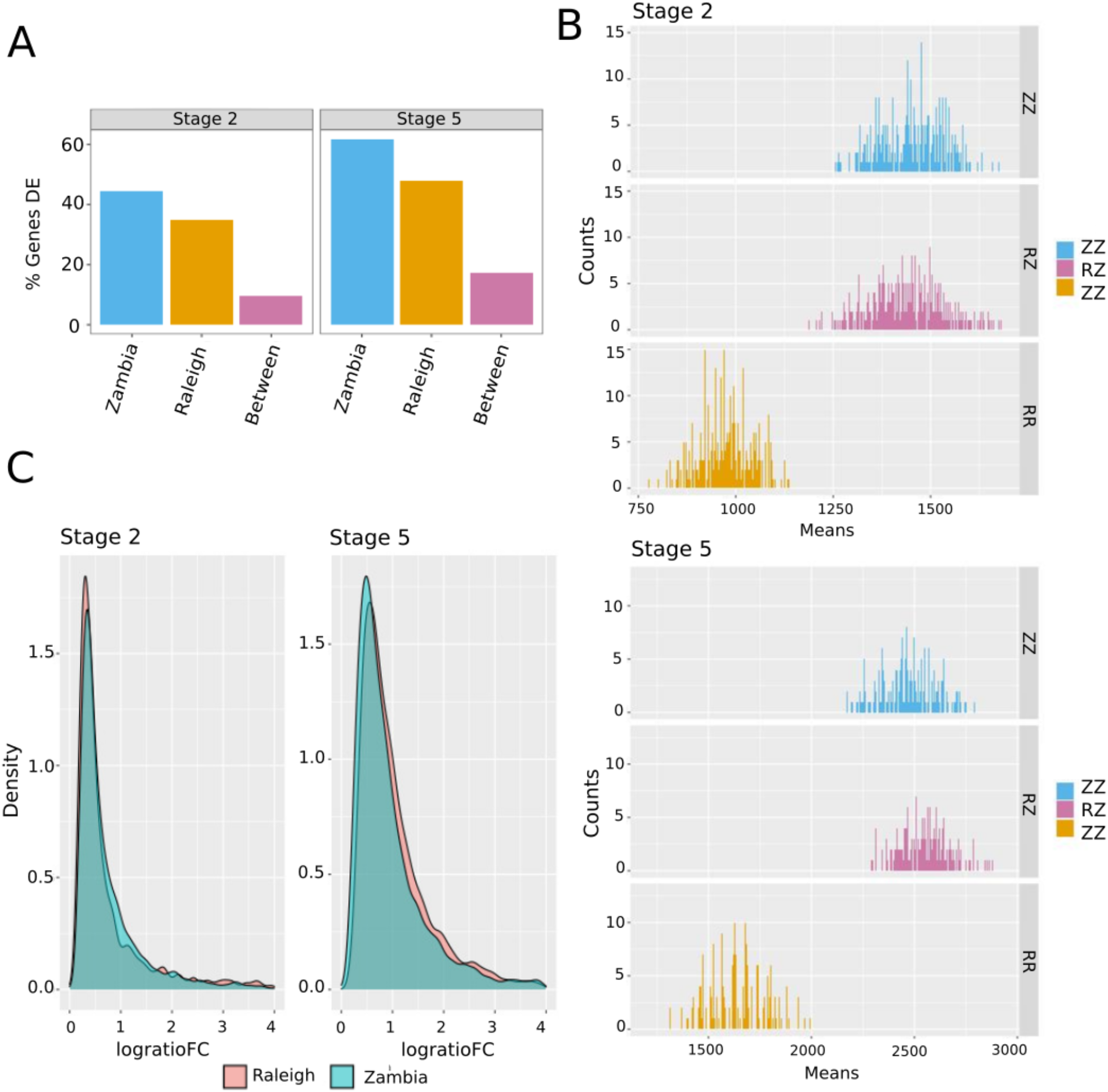
Differential expression within and between populations. (Higher number of differential expression within Zambia. Larger magnitude of changes within Raleigh at Stage 5) A) Percent of genes differentially expressed within and between the Zambia and Raleigh populations at stage 2 and stage 5. More differences are found within populations (blue, orange) than between populations (pink). (B) To control for the number of comparisons within and between lines, we also examined pairwise differences between lines at each stage (stage 2 top, stage 5 bottom). When compared in this way, at both stages, the distributions of DE genes within the Zambia population and between the Zambia and Raleigh populations are similar, with fewer DE genes within the Raleigh lines. (C) Distributions of the magnitudes of differences in expression in DE genes, which shows that the magnitude of changes between differentially expressed genes is greater within the Raleigh population at stage 5 than the Zambia population at this stage.

We asked if there were similarities in the identity of genes with differential expression within populations at the two stages. A proportion of genes were found to be differentially expressed within both populations at stage 2 and stage 5 (Figure 2A). Of all the DE genes at stage 2 combined, 43% were only DE within Zambia and 28% within only Raleigh lines, while 29% of genes were DE in both (Supplemental Figure S1). At stage 5 the percent of genes only DE within the Zambia lines stayed relatively similar at 39% whereas the percentage of genes only varying expression within the Raleigh population was lower at 20%, due to a higher proportion of genes in both (at 41%; Supplemental Figure S1). Thus, the percentage of genes varying in expression levels in both populations is higher in stage 5 than stage 2. Therefore, there is a common set of genes that vary in transcript levels within both populations in addition to a unique set of genes that vary only within the respective populations, and these vary by stage, with more shared differences at stage 5.

### Differences in the Magnitude of Expression Variation Within Populations

With more genes differentially expressed within the Zambia population than the Raleigh population, we asked if the magnitude of expression changes were similar between populations. To do this, we found the maximum and minimum expression value for each differentially expressed gene within the populations. From this, we computed the log ratio of the fold change for each DE gene. We then asked if the distribution of the log ratio of fold changes for DE genes were different between the two populations at either stage (Figure 2C). There is no significant difference between the means of log ratio of fold changes when comparing stage 2 between populations (t-test, p = 0.9109), thus no evidence that the magnitude of transcript abundance changes is different between populations. There is, however, a significant difference between the means of the log ratio of fold changes between the two populations at stage five (t-test, p= 7.278e-06) with a higher magnitude of fold changes within the Raleigh population. Therefore, although there are fewer genes differentially expressed within the Raleigh population at stage 5, the magnitude of these differences are on average higher than the genes differentially expressed within the Zambia population at this stage.

### More Differences within Populations Than Between Populations at Maternal and Zygotic Stages

Next, we asked if there were fixed expression differences between the populations. We define fixed expression differences as genes that are on average higher, or lower, in one population than the other (i.e. have similar levels in all lines from a population, that are significantly different than all the lines in the other population; see Figure 3A for examples). We used the Raleigh lines and the Zambia lines as replicates in DE analysis. Similar to the expression variation within populations, the percentage of genes that were differentially expressed between populations increased from stage 2 to stage 5 (Figure 2A). We find that there are more genes differentially expressed within populations than fixed expression differences between the populations at both stages (Figure 2A).

**Figure 3.**
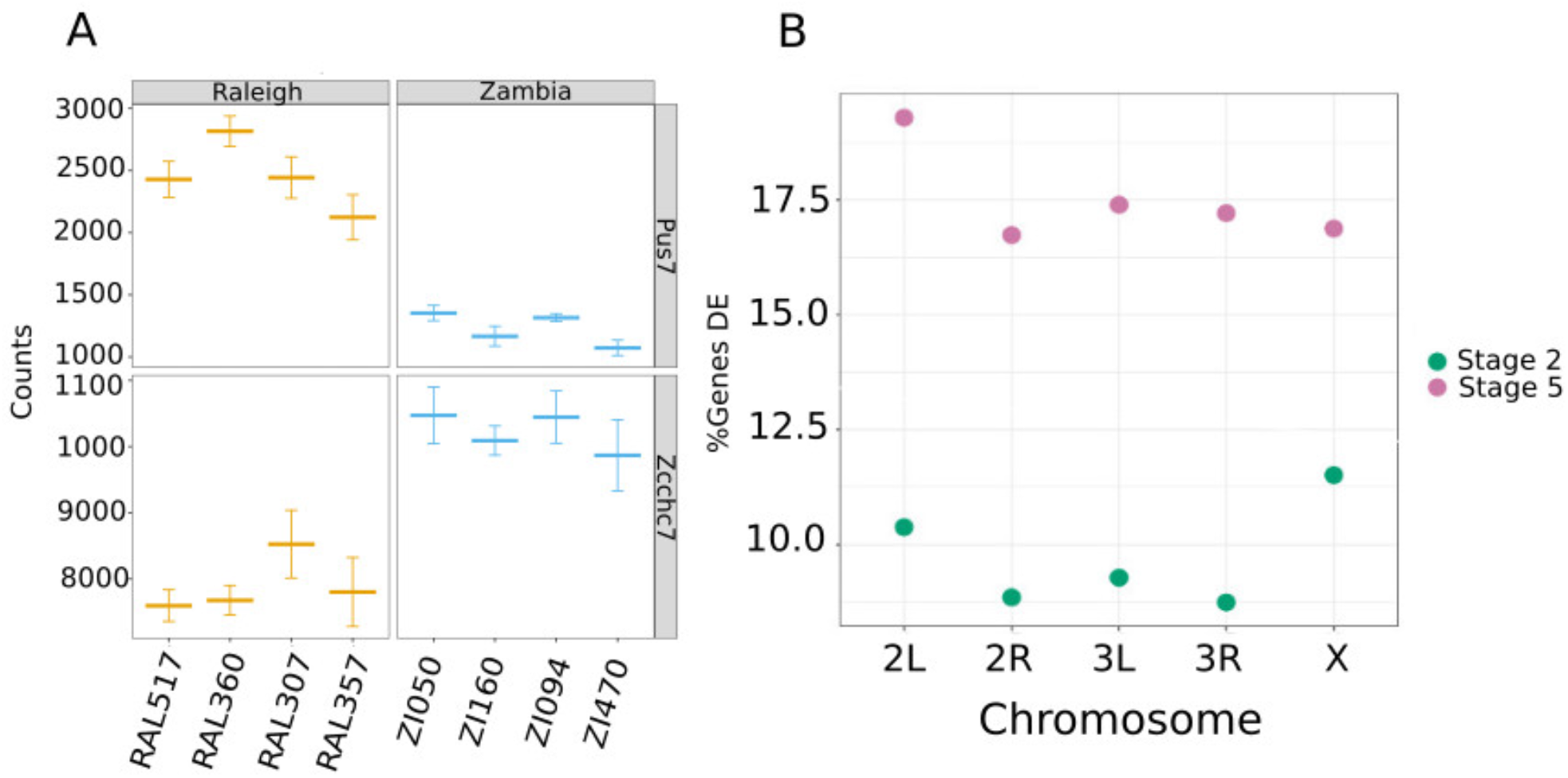
Examination of putative fixed differences between populations. A) Expression levels in counts for two example genes, showing what we categorize as fixed differences in transcript levels between populations. B) Percentage of genes that are differentially expressed as compared to the number of genes on the chromosome at each stage. At stage 2, where all transcripts are maternal in origin, there is a significant enrichment of DE genes on the X chromosome.

In addition to finding fixed expression differences, we asked how many genes were differentially expressed between individual lines. Genes differentially expressed between lines from different populations in the pairwise analysis represent differences only between the two lines in the comparison, rather than fixed expression differences between the two populations as in the previous analysis. This resulted in DE analysis between every pair of lines resulting in 28 of total comparisons.12 DE tests between lines of the same population (RR and ZZ), and 16 DE tests between lines of different populations (RZ). Since there are fewer tests between lines of the same population than between lines of different populations we used bootstrapping in order to compare the average number of DE genes between these categories. Similarly to the previous within population analysis, there are fewer DE genes between individual Raleigh lines (RR) than Zambia lines (ZZ) at both stages (Figure 2B). Interestingly, we find that the average pairwise differences between lines (RZ) of different populations at stage 2 was not significantly [p = 0.06972; Wilcoxon rank sum test] different than the average pairwise differences between Zambia lines (ZZ) at this stage (Figure 2B). However, at stage 5, the average number of differences between lines of different populations are higher relative to the number of differences between Zambia lines [p < 2.2e-16; Wilcoxon rank sum test]. Therefore, there is as much variation of expression between individual Zambia lines at stage 2 as between individual lines from different populations at this stage. In contrast, variation between individual lines from different populations at stage 5 surpasses the differences between individual Zambia lines at this stage.

### More Expression Variation Between than Within Species

Expanding our analysis, we investigated gene expression variation within and between species of *Drosophila* at maternal and zygotic stages. In a previous study, we generated RNA-seq data from *D. simulans, D. sechellia, D. yakuba* and *D.erecta* from stage 2 and stage 5 embryos using the same single embryo RNA extraction method implemented here. We chose these two pairs of sister species as they are closely related, but one pair (*D. simulans* and *D. sechellia*) diverged more recently (~250,000 years ago, (McDermott & Kliman, 2008) than the other pair (*D. yakuba* and *D. erecta*, estimated 8 MYA divergence time, (see TimeTree). RNAseq reads from these species were processed identically to the *D. melanogaster* reads for this analysis (see Methods). Genes considered in this analysis were limited to one-to-one orthologs across the 5 species, a total of 12,110 genes. As we had only one line per the other species, we performed the DE analysis pairwise for each of our *D. melanogaster* lines, as well as between each pair of sister species. Number of DE genes in each population or species were normalized to the number of genes transcribed at each stage to compare the percentage of DE genes at both stages and across species. From these comparisons, within and between species there are more differentially expressed genes at stage 5 than stage 2 (Figure 4). For maternal genes, the more closely related species pair *D. simulans* and *D. sechellia* have the highest proportion of genes DE. While most *D. melanogaster* lines have fewer differences than either of the species comparisons at this stage, two of the Raleigh vs. Zambia comparisons have as high of a proportion of their maternal genome maternally expressed as the more distantly related species pair, *D. yakuba* and *D. erecta*. For stage 5, both species pairs have a larger proportion of their transcripts DE than any of the within-species comparisons. In all, both stages have, on average, fewer genes DE for within-species comparisons than between species, but this pattern is much stronger for stage 5, a stage with more genes DE in all comparisons.

**Figure 4.**
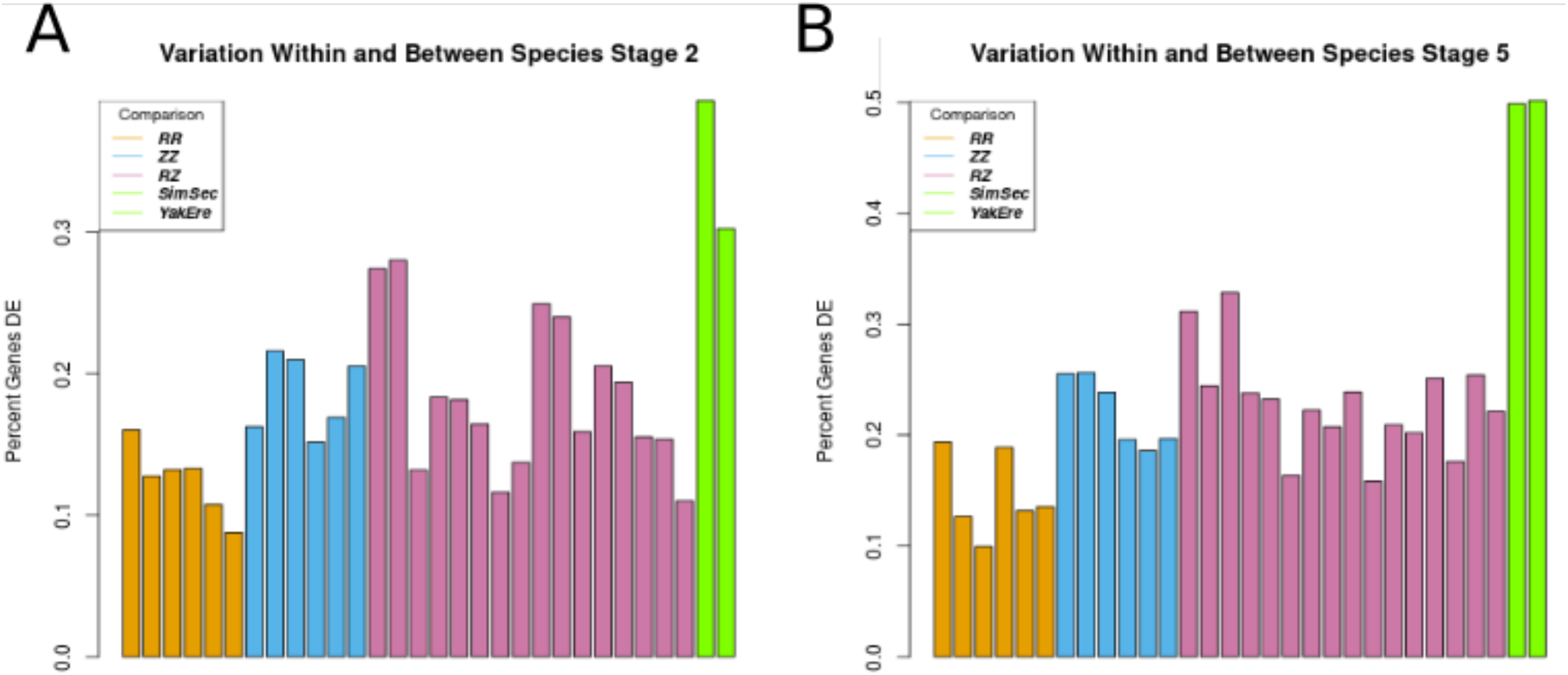
Fewer differences in differentially expressed genes within species than between species. DE analysis was done between pairs of Raleigh lines (orange), pairs of Zambia lines (blue), between lines of the two populations (purple) and between species pairs (green; representing one line from each species) from stage 2 and stage 5 embryos (see Methods). The between species DE analysis was done with pairs of closely related species: *D. simulans* and *D. sechellia* (SimSec) as well as *D. yakuba* and *D.erecta* (YakEre). It was found that there were on average fewer DE genes between lines of *D. melanogaster* than between pairs of species at both stages. Stage 5 has a higher proportion of genes identified as DE for each comparison, and the between-species comparison has an even higher enrichment in DE genes as compared to stage 2, where more constraint may be present.

### Enrichment of DE Genes at Maternal Stage on the X Chromosome

Before the zygotic genome is activated, embryonic development is entirely under control of maternal gene products. Therefore all stage 2 transcriptomes are supplied entirely by XX genomes and the zygotic genome is transcribed by either XX or XY genomes. Given the possibility of different evolutionary pressures, we asked whether there is a difference of enrichment of DE genes on the autosomes or X chromosome across maternal and zygotic stages. In *Drosophila*, dosage compensation occurs when genes on the X in males have increased transcription to maintain equal levels of gene products to females (Lucchesi & Kuroda 2015; Conrad & Akhtar 2012). This occurs sometime after stage 5 (Georgiev et al., 2011). However, all stage 5 embryos collected are male and are therefore directly comparable at this stage. We normalized the number of DE genes per chromosome by the number of genes expressed on each chromosome.

Interestingly, we found that DE genes at stage 2 between populations were enriched on the X chromosome compared to the autosomes (Figure 3B). However, enrichment of DE genes on the X chromosome is absent at stage 5. Maternal transcripts are not completely degraded by stage 5, so we also asked if the trend seen for all of stage 5 transcripts were the same for transcripts that are zygotic only. As expected, fixed expression differences between zygotic only genes were not enriched on the X chromosome (Fisher’s exact test, p <0.05) having the same result as all genes at stage 5.

### A number of the most differentially expressed genes are genes with known selection signatures

A number of the most differentially deposited transcripts between populations are genes that have been shown previously to have signatures of selection at the level of the genome under different conditions. For example, a previous study found that genes within the chemosensory system have undergone local adaptation following *D.melanogaster*’s global expansion out of Africa (Arguello et al., 2016). This study was based on the genomes of five different geographically distinct populations of *D. melanogaster* including both North American and African populations. Notable within the top ten most DE maternally deposited genes between populations is Gstd9, a glutothione-S-transferase, which belongs to a gene family that was found to have signals of selective sweeps upon global expansion (Arguello et al., 2016). In total, seven glutothione-S-transferases were found to be differentially deposited between the Raleigh and Zambia populations. In the same study (Arguello et al., 2016) the zinc finger protein family was shown to have strong population differentiation. Zcchc7, a zinc-finger protein, is also among the top ten most differentially deposited transcripts. These two genes both have undergone dramatic qualitative changes in maternal deposition (Figure 3A and Figure 6A).

The second most significantly differentially deposited transcript is the actin binding protein Unc-115a. The paralog of this gene, Unc-115b, was also found to be differentially expressed between populations. Both these genes have higher expression levels in the Raleigh population. Interestingly, Unc-115b was found in a previous study to be the most highly upregulated gene in a strain resistant to the insecticide DDT 91-R compared to a DDT compromised strain, 91-C (Seong et al., 2017). Unc-11b was one of two genes found in this study to be highly upregulated across all stages of development that were assayed (Seong et al., 2017). This gene was found to be in one of six selective sweeps that coincided with constitutive expression differences between DDT resistant and compromised lines.

### A number of the most differentially expressed genes are annotated as pseudogenes

The most differentially maternally deposited gene between the Zambia and Raleigh populations in our analysis is the gene CR40354 which is annotated in the *D.melanogaster* genome as a pseudogene attribute with unknown function. This prompted us to investigate other genes annotated as pseudogenes in our dataset, especially given that previous annotations that identified these genes as pseudogenes were more likely to have been done in non-African populations. We asked how many pseudogenes were maternally deposited and zygotically expressed within and between populations. A total of 69 and 70 genes labeled as pseudogenes were found to be maternally deposited within the Raleigh and Zambia populations, respectively. A total of 46 and 49 genes labeled as pseudogenes were found to be expressed zygotically only in the Raleigh and Zambia populations. Between the populations, 18 pseudogenes were found to be differentially maternally deposited. One pseudogene which caught our attention was the *swallow Ψ (swaΨ*) pseudogene which is differentially expressed within the Zambia population in our analysis. *swaΨ* is a result of a recent genome duplication of swallow, and is only found in *D. melanogaster* (Chao et al., 1991). *Swallow* is a critical gene to early development, and is required for proper Bicoid positioning in the embryo (Stephenson et al., 1988). Previous studies (Petrov et al., 1998) have suggested that *swaΨ* not transcribed in *D. melanogaster*. We found it to be very lowly expressed in the Raleigh lines, but variably expressed within the Zambia lines with one line, ZI160, showing relatively high expression levels (Figure 5D). To investigate further, we sequenced the swaΨ locus in each of the lines. We found very few single nucleotide variants in this gene, however we also discovered a 15bp population-specific deletion. All Raleigh lines have a 15bp deletion in the annotated exon 3 of swaΨ, which is present in all four Zambian lines. This sequence is part of the fully functional exon 3 of the swallow gene.

**Figure 5.**
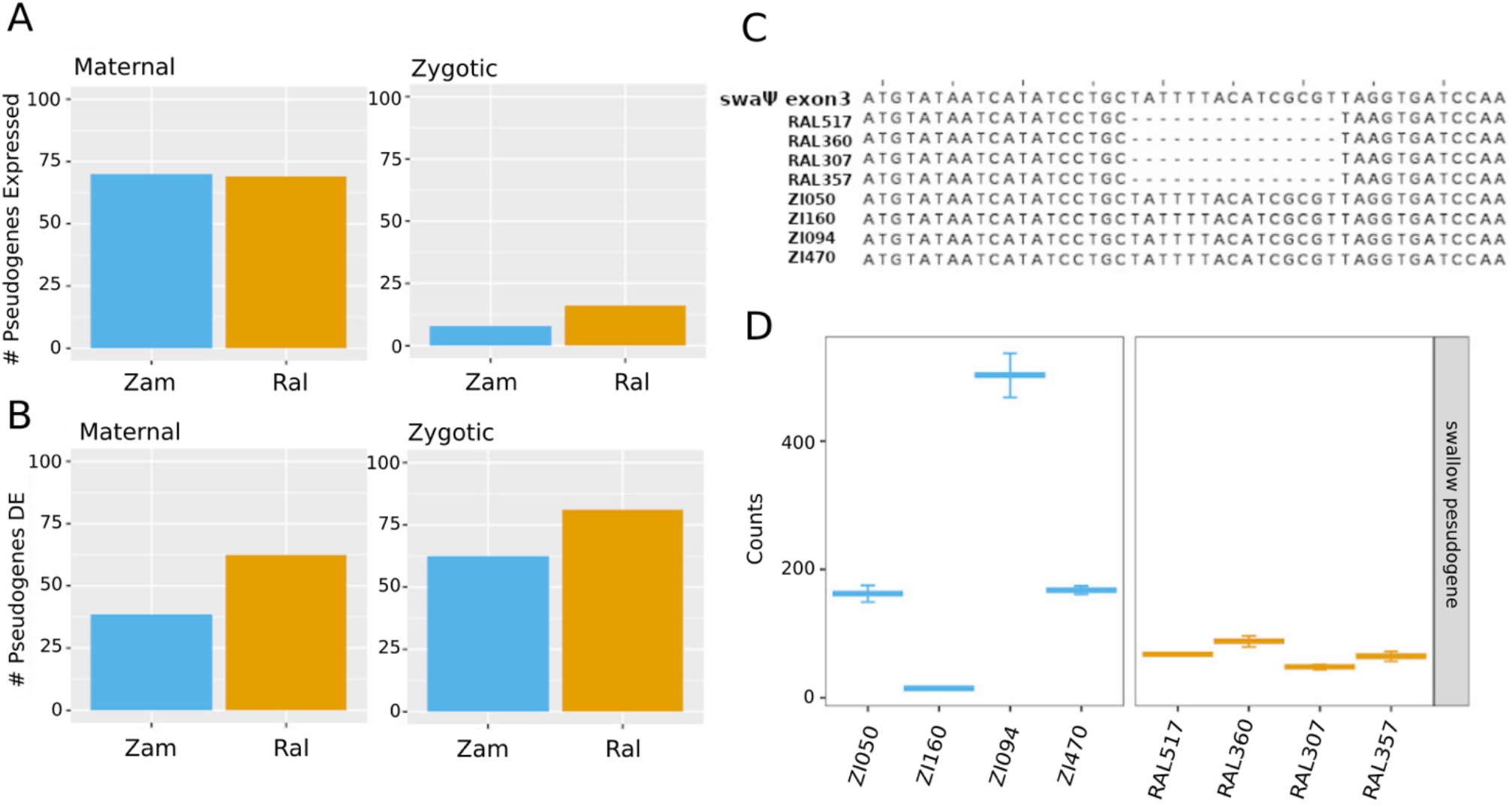
Variation in pseudogene transcript levels. A) At each stage (maternal, zygotic), the number of annotated pseudogenes expressed are similar between populations. The smaller number of pseudogenes expressed at the zygotic stage reflect that this analysis was restricted to zygotic-only genes, which are zygotic genes with no maternal expression. B) Of the pseudogenes expressed at each stage, a larger proportion are differentially expressed in the Raleigh lines. C) One example, the *swallow* pseudogene, has a 15bp deletion shared by all of the Raleigh lines at the position shown in the alignment. D) The *swallow* pseudogene is more highly expressed in a number of the Zambia lines, with considerable variation between lines.

### Variation in Heat shock Proteins

Modifying maternal RNAs and proteins in the embryo can have effects on development, phenotypes and ultimately fitness (Driever & Nüsslein-Volhard 1988, Zhang et al. 1998). One gene family that is critical to survival are heat shock proteins (Feder & Hofmann 1999, Kregal 2001). In total, 34 heat shock proteins were found to be differentially deposited within the Raleigh population and 32 heat shock proteins were differentially deposited within the Zambia population. This is in contrast to after zygotic genome activation, where no zygotic only heat shock proteins were found to be differentially expressed within the two populations. Previous work by Lockwood et al. has shown evidence that increased maternal deposition of a heat shock protein increases embryo thermal tolerance in *D. melanogaster* (Lockwood et al., 2017). Interestingly, Hsp23 was found to be differentially deposited in the lines that we examined (Figure 6A, bottom panel). Specifically, the levels of Hsp23 mRNA in ZI094 is between 4-14X higher than the other three Zambia lines and 11-600X higher levels than the Raleigh lines, all which have variable expression. This overall trend persists at stage 5, with mean levels of Hsp23 increasing in ZI094 and maintaining higher expression levels compared to all other lines. Based on this observation, we performed heat shock experiments on all of the lines to assay differences in embryo survival after heat stress (see Methods). In general, we found that heat shock tolerance does not correspond in a predictive way with levels of heat shock transcripts (Figure 6B).

**Figure 6.**
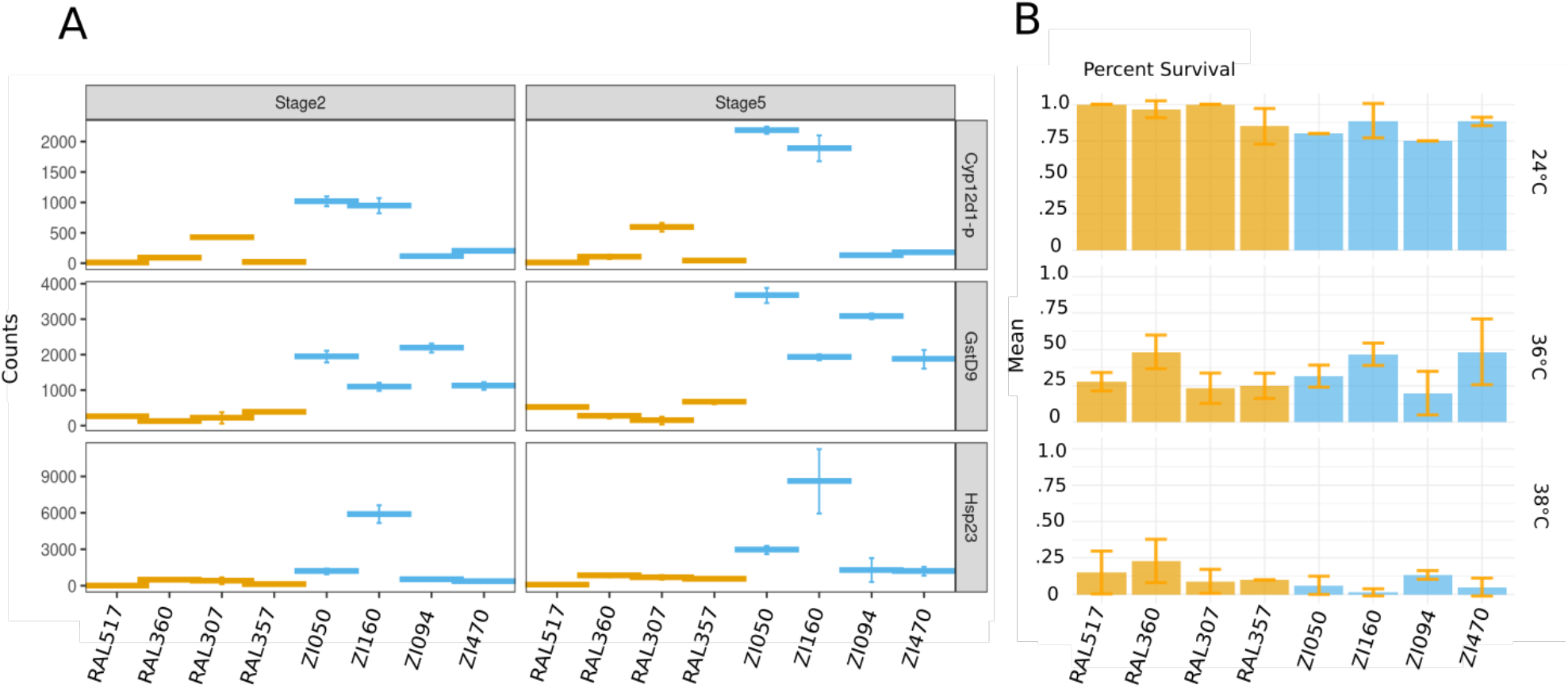
Examples of differentially expressed genes with previous evidence for functional significance. A) Transcript levels for three example genes, shown at both developmental stages labeled across the top, for the Raleigh lines (blue) and the Zambia lines (orange). B) Results of experiments testing survival of embryonic heat shock across lines, showing relative survival at three temperatures. While on average the Raleigh lines have higher survival after heat shock at 24°C and 38°C, they also have higher survival at standard rearing temperatures, results do not correspond well with heat shock transcript levels.

## Discussion

While previous studies have shown that the maternal and early zygotic transcriptomes are highly conserved across species (Atallah & Lott 2018, Preuss et al. 2012; Heyn et al. 2014), here we show that that there is variation present in gene expression on the shorter evolutionary timescale represented within a species, *D. melanogaster*. We chose lines from Zambia and Raleigh, North Carolina to encompass a broad span of genetic diversity within and among populations.

Our results show that the transcriptomic dynamics at these developmental stages reflect what is known about the population genetic history of *D. melanogaster* from genomic studies. Previous studies found more genetic variation within African populations than non-African populations (Langley et al. 2012, Begun & Aquadro 1993, Pool & Aquadro 2006, Baudry, Viginier, & Veuille 2004) we found the same pattern with the maternal and early zygotic transcriptomes. There are differential transcript abundances within both the Zambia and Raleigh populations, and some of the same transcripts are variable in each population, but there is more population-specific variation within the Zambia lines. We also find that with pairwise comparisons between lines, the Raleigh lines have far fewer genes identified as differentially expressed, but comparisons within Zambia have as many (stage 2) or only slightly fewer (stage 5) differentially expressed genes as when comparing lines from the two populations. The increased number of differentially expressed genes in the Zambia lines is consistent with high levels of genomic variation found in the ancestral range of this species (Baudry, Viginier, and Veuille 2004), while reduced number of differentially expressed genes in Raleigh likely reflects the lower genetic polymorphism levels following the out-of-Africa bottleneck (David & Capy 1988,Begun & Aquadro 1993). Interestingly, while consistent with the genomic variation within these lines, our results stand in contrast to microarray studies in adults which found less transcript variation within African and non-African populations than between, which has been taken as a sign of directional selection (Müller et al. 2011, Hutter et al. 2008).

Also consistent with previous genomic studies are the numbers of genes highlighted by our DE analysis that have also been identified in studies performing artificial selection or population genomic studies on the global expansion of the species (Arguello et al. 2016, Seong et al. 2017). Many are used as examples throughout the manuscript, but many have been associated with xenobiotic metabolism (GstD9, Cyp12d1-p), possible environmental adaptation (Zcch7), and DDT resistance (Unc-115a, Unc-115b). Thus, many of our most significantly DE genes are also likely under selection, and their functions are consistent with adaptation to a new environment. Studies to determine the adaptive function of these genes are often carried out in adults (Yang et al. 2007, Strycharz et al. 2013), our data suggests that these differences in transcript level are also present in the embryo, and thus may potentially be of adaptive value at this stage.

Previous studies have found an especially high degree of conservation of the maternal transcriptome across species (Atallah & Lott 2018, Heyn et al. 2014, Preuss et al. 2012), this study provides evidence this is also true within *D. melanogaster*. Whether examining the number or proportion of differentially expressed genes within populations, between populations, between pairs of lines, or between species, there are fewer differences in transcript level found at stage 2 when all transcripts are maternal than at stage 5 after zygotic genome activation. The analysis of proportions of genes DE within and between species is especially suggestive relative to these stage-specific dynamics. At stage 5, the proportion of genes DE between species is far higher than the within-*D. melanogaster* comparisons, as well as showing a higher proportion of DE genes overall in every comparison. In contrast, at stage 2, there are fewer genes DE in each comparison, and the between species comparisons (while still higher on average than the within-*D. melanogaster* comparisons) are only slightly higher. This suggests that relative to one another, more of the maternal transcriptome may be under stabilizing selection than the more rapidly evolving zygotic stage transcriptome (Nuzhdin et al., 2004).

In conclusion, we find that while the maternal and zygotic transcriptomes, while generally conserved, do show some interesting differences in transcript abundance even in the relatively short period of evolutionary time represented by the diversity within a species. This species, *D. melanogaster*, has more variation for transcript abundance at these critical developmental stages within populations than between them. And consistent with what has been determined between Drosophila species (Atallah & Lott 2018), we show that the maternal transcriptome is more highly conserved than the zygotic transcriptome, and more of the maternal genome may be under purifying selection. Together, the presented data highlight how a constrained developmental trait evolves over short periods of evolutionary time.

---

**Supplemental Figure S1.**
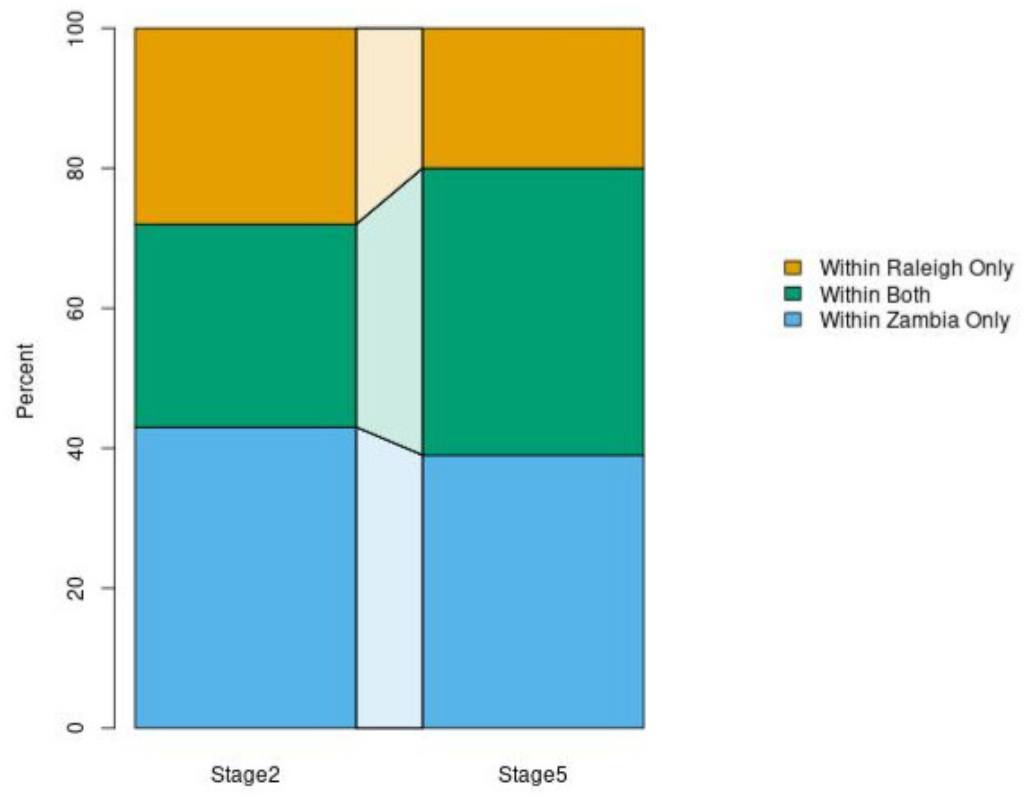
Shared expression variation within populations. At both stage 2 and stage 5 there are a set of genes that are differentially expressed within both the Raleigh and Zambia populations (green). This shared set of genes differentially expressed in both populations increasing from stage 2 to stage 5.

